# Comparing FRET pairs that report on intersubunit rotation in bacterial ribosomes

**DOI:** 10.1101/2023.05.09.540051

**Authors:** Ananya Das, Chen Bao, Dmitri N. Ermolenko

## Abstract

Mediated by elongation factor G (EF-G), ribosome translocation along mRNA is accompanied by rotational movement between ribosomal subunits. Here, we reassess whether the intersubunit rotation requires GTP hydrolysis by EF-G or can occur spontaneously. To that end, we employ two independent FRET assays, which are based on labeling either ribosomal proteins (bS6 and bL9) or rRNAs (h44 of 16S and H101 of 23S rRNA). Both FRET pairs reveal three FRET states, corresponding to the non-rotated, rotated and semi-rotated conformations of the ribosome. Both FRET assays show that in the absence of EF-G, pre-translocation ribosomes containing deacylated P-site tRNA undergo spontaneous intersubunit rotations between non-rotated and rotated conformations. While the two FRET pairs exhibit largely similar behavior, they substantially differ in the fraction of ribosomes showing spontaneous fluctuations. Nevertheless, instead of being an invariable intrinsic property of each FRET pair, the fraction of spontaneously fluctuating molecules changes in both FRET assays depending on experimental conditions. Our results underscore importance of using multiple FRET pairs in studies of ribosome dynamics and highlight the role of thermally-driven large-scale ribosome rearrangements in translation.

## Introduction

During the elongation cycle, following the formation of a new peptide bond, the ribosome translocates along mRNA by one codon in the process mediated by universally conserved elongation factor G (EF-G or EF-2 in eukaryotes). In the course of translocation, peptidyl- and deacylated tRNAs move from A and P into P and E sites of the ribosomes, respectively. Translocation is accompanied by a cycle of forward and reverse rotation between the small and large ribosomal subunits [1, 2]. Early chemical probing studies revealed that peptide bond formation causes resulting peptidyl- and deacylated tRNAs to spontaneously translocate on the large subunit into P and E sites, respectively, while their anti-codon stem-loops remain in the original A and P sites on the small subunit [3]. Thus, tRNAs spontaneously adopt hybrid A/P and P/E conformations [3]. Structural and FRET studies indicated that hybrid state formation is accompanied by ∼8-10° rotation of the small 30S subunit relative to the large 50S subunit [4–8]. EF-G induces translocation of mRNA and tRNAs on the small subunit coupled to reverse rotation between subunits into the non-rotated conformation, in which tRNAs adopt P/P and E/E states [9–11]. Further structural studies revealed additional intermediates and rearrangements of the ribosome [8]. Nevertheless, translocation can generally be described as transition from the non-rotated (NR) to rotated (R) and then back to non-rotated (NR) conformations.

Two popular single-molecule (sm)FRET assays, which involve labeling either ribosomal proteins (bS6 and bL9) [9, 12] or rRNA (h44 of 16S and H101 of 23S rRNA) [11], have been extensively used to follow intersubunit rotation in bacterial ribosomes. Protein labeling involves introducing unique cysteine residues into proteins bS6 and bL9 for fluorescent labeling followed by incorporation of fluorescently-derivatized proteins bS6 and bL9 into ΔbS6 30S and ΔbL9 ribosomes, respectively [9, 13]. To label rRNA, 23 nucleotide-long extensions are engineered into the loops of helixes h44 of 16S and H101 of 23 rRNA, respectively [11]. These extensions are then hybridized with fluorescently-labeled DNA oligos. Both S6/L9 and h44/H101 smFRET assays detect the NR→R→NR transition during the elongation cycle [14, 15]. Both FRET assays also indicated that NR→R is triggered by peptidyl-transferase reaction, which quickly follows A-site tRNA binding [9, 11, 12]. S6/L9 and h44/H101 smFRET assays, however, produced conflicting results regarding the R→NR transition.

The S6/L9 FRET pair revealed that in the absence of EF-G, pre-translocation ribosomes containing deacylated P-site tRNA undergo repetitive spontaneous fluctuations between the R and NR conformations that do not result in mRNA translocation [12]. Intersubunit dynamics observed in S6/L9 FRET experiments was similar to spontaneous tRNA fluctuations between the classical (A/A and P/P) and hybrid (A/P and P/E) states [16–19] and inward-outward movements of the 50S L1 stalk [18, 20–22], which were detected by smFRET in pre-translocation ribosomes in the absence of EF-G. These findings indicated coupling between intersubunit rotation, movements of the 50S L1 stalk and tRNA transitions between hybrid and classical states. Indeed, structural data demonstrated that the hybrid and classical states of tRNA binding were exclusively observed in the R and NR conformations of the ribosome, respectively [8, 23]. Taken together with tRNA and L1 stalk fluctuations in pre-translocation ribosomes, spontaneous intersubunit rotation detected by S6/L9 FRET pair was interpreted as a key supporting evidence for the model suggesting that EF-G•GTP acts by converting thermally-driven conformational dynamics of the ribosome and tRNAs into ribosome translocation along mRNA [2, 7, 24].

In contrast to smFRET data obtained using S6/L9 FRET pair, h44/H101 smFRET assay showed that in the absence of EF-G•GTP, pre-translocation ribosomes remain idle in the R state [11]. The R to NR transition in h44/H101-labeled ribosomes was strictly dependent on EF-G•GTP binding [11]. Experiments with h44/H101 FRET pair led to the model suggesting that energies of peptidyl transfer and GTP hydrolysis by EF-G are used to surmount activation barriers to forward and reverse intersubunit rotation, respectively [11, 25]. Since in the absence of EF-G, tRNAs were shown to spontaneously fluctuate between classical and hybrid states [16, 19], h44/H101 FRET data also implied that tRNA fluctuations occurred in the R conformation of the ribosome and thus were uncoupled from intersubunit rotation.

To reconcile differences between S6/L9 and h44/H101 smFRET data, it was hypothesized that the S6/L9 assay reports on a rearrangement of the ribosome, which is different from intersubunit rotation, e.g. 50S L1 stalk movement [11] or 30S head rotation [26]. Another possibility is that differences in experimental conditions, such as ion and polyamine concentrations, may underlie discrepancies between S6/L9 and h44/H101 smFRET data. Finally, extending rRNA stem-loops for fluorescent labeling in h44/H101 construct might perturb structural dynamics of the ribosome [2]. Molecular dynamics simulations indicated that even in the absence of such labeling-induced perturbations, the intrinsic structural flexibility of the ribosome can complicate FRET detection of intersubunit rotation [27]. While changes in the distance between sites of labeling in both S6/L9 and h44/H101 pairs were shown to correlate with intersubunit rotation, inter-fluorophore distances in the h44/H101 pair consistent with the R conformation could be observed with some frequency in the NR conformation of the ribosome. These molecular dynamics simulation data also showed that S6/L9 FRET pair is more responsive to intersubunit rotation [27].

To test these hypotheses, we perform side-by-side comparison of S6/L9 and h44/H101 FRET pairs under the same experimental conditions. Our data indicate that both FRET pairs detect intersubunit rotation. Both FRET pairs also reveal spontaneous repetitive intersubunit rotation in pre-translocation ribosomes in the absence of EF-G. However, significantly fewer h44/H101-labeled ribosomes undergo spontaneous fluctuations than S6/L9 ribosomes. The difference in dynamics between S6/L9 and h44/H101 FRET pairs can be further amplified by variations in buffer conditions. Our data highlight importance of employing several complementary FRET pairs in studies of structural dynamics of large macromolecular machines such as the ribosome.

## Results

### S6/L9 and h44/H101 FRET pairs detect NR, R and semi-R conformations of the ribosome

To test whether S6/L9 and h44/H101 FRET pairs truly report on intersubunit rotation in the ribosome, we measured FRET between a donor (Cy3) and an acceptor (Cy5) fluorophores in ribosome complexes, which represent different steps of translation initiation and elongation. In S6-Cy5/L9-Cy3 ribosome, we observed ∼0.6 and ∼0.4 FRET, which were previously attributed to the NR and R conformations, respectively (Fig. 1-2). In h44-Cy3/H101-Cy5 ribosomes, we observed ∼0.4 and 0.2 FRET, which likely also correspond to the NR and R conformations, respectively (Fig. 1-3). These values were somewhat lower than 0.45 and 0.35 FRET values previously reported for h44-Cy3/H101-Cy5 ribosomes. The shift to lower FRET values in our experiments is likely due to differences in microscope fluorescence detection (i.e. emission filters and the dichroic mirror).

**Figure 1.**
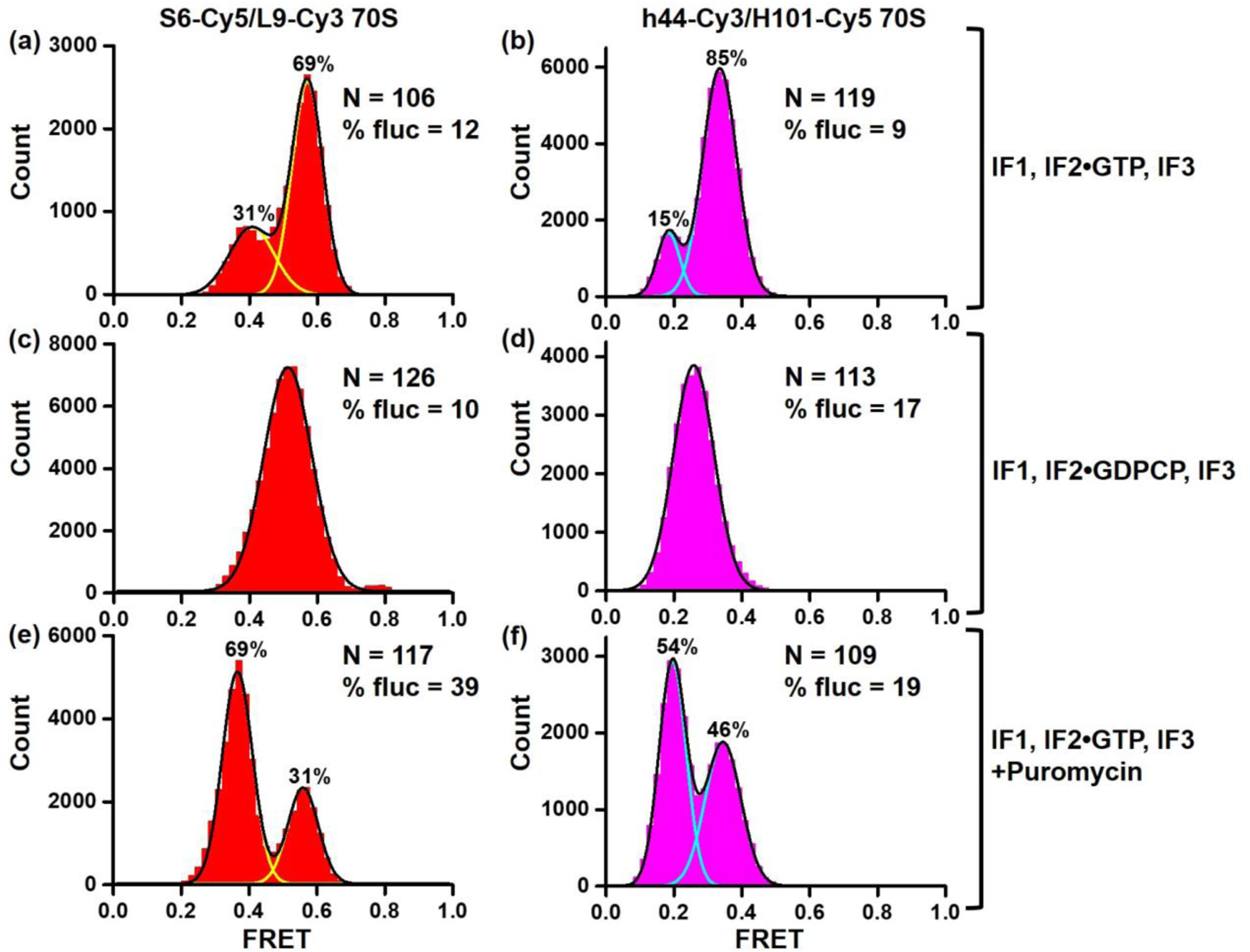
Detecting NR, R and semi-R confromations of the ribosome using S6-Cy5/L9-Cy3 and h44-Cy3/H101-Cy5 FRET assays. Histograms show FRET distribution in S6-Cy5/L9-Cy3 (a, c, e) and h44-Cy3/H101-Cy5 (b, d, f) ribosomes imaged in polyamine buffer. 70S ribosomes containing P-site fMet-tRNA^fMet^ were assembled in the presence of IF1, IF2•GTP, IF3 (a-b, e-f) or IF1, IF2•GDPCP, IF3 (c-d). N indicates the total number of FRET traces compiled into each histogram. % fluc designates the percentage of traces that show fluctuations. Yellow and cyan lines indicate individual Gaussian fits. Black line shows the sum of Gaussian fits. The fraction of the ribosomes in the R and NR conformations are displayed above the corresponding Gaussian peaks.

**Figure 2.**
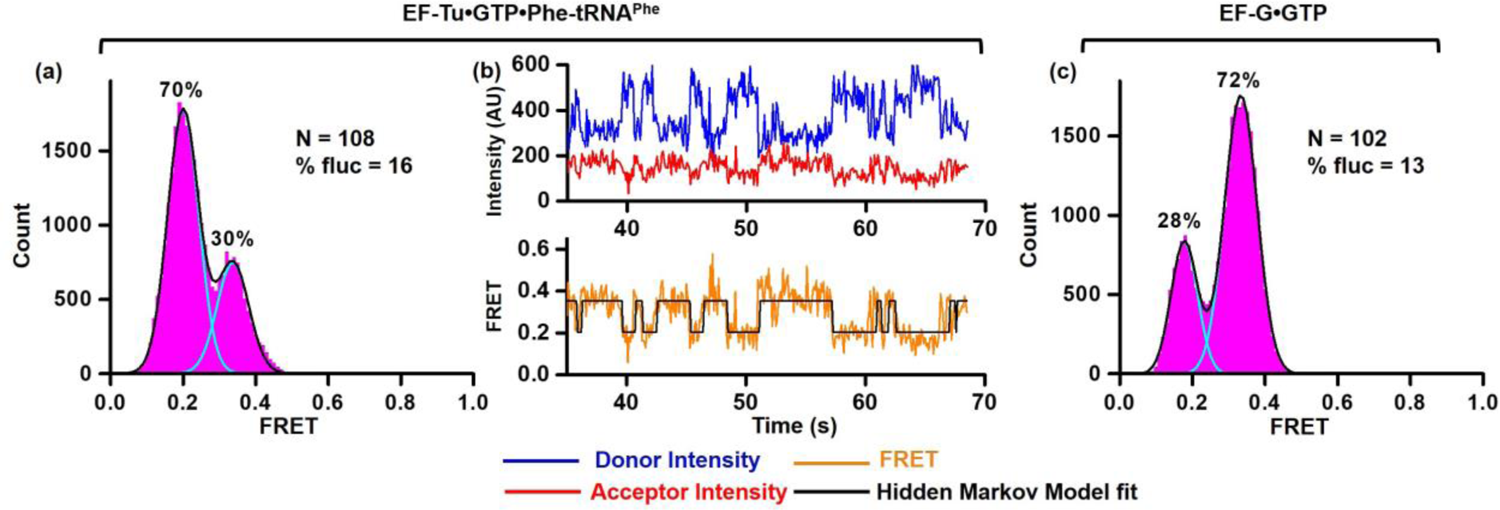
Spontaneous intersubunit rotation in pre-translocation h44-Cy3/H101-Cy5 ribosomes. h44-Cy3/H101-Cy5 ribosomes bound with P-site fMet-tRNA^fMet^ were first incubated with EF-Tu•GTP•Phe-tRNA^Phe^ (a, b) and then with EF-G•GTP (c). (a, c) Histograms show FRET distribution. N corresponds to the total number of FRET traces compiled into each histogram. % fluc indicates the percentage of traces that exhibit FRET fluctuations. Cyan and black lines show individual Gaussian fits and the sum of Gaussian fits, respectively. The fraction of the ribosomes in rotated and non-rotated conformations are shown above the corresponding Gaussian peaks. (b) smFRET trace shows spontaneous intersubunit rotation in h44-Cy3/H101-Cy5 pre-translocation ribosomes. Fluorescence intensities of Cy3 (donor) and Cy5 (acceptor) are shown in blue and red, respectively. FRET efficiency and HMM fit are displayed in orange and black, respectively.

**Figure 3.**
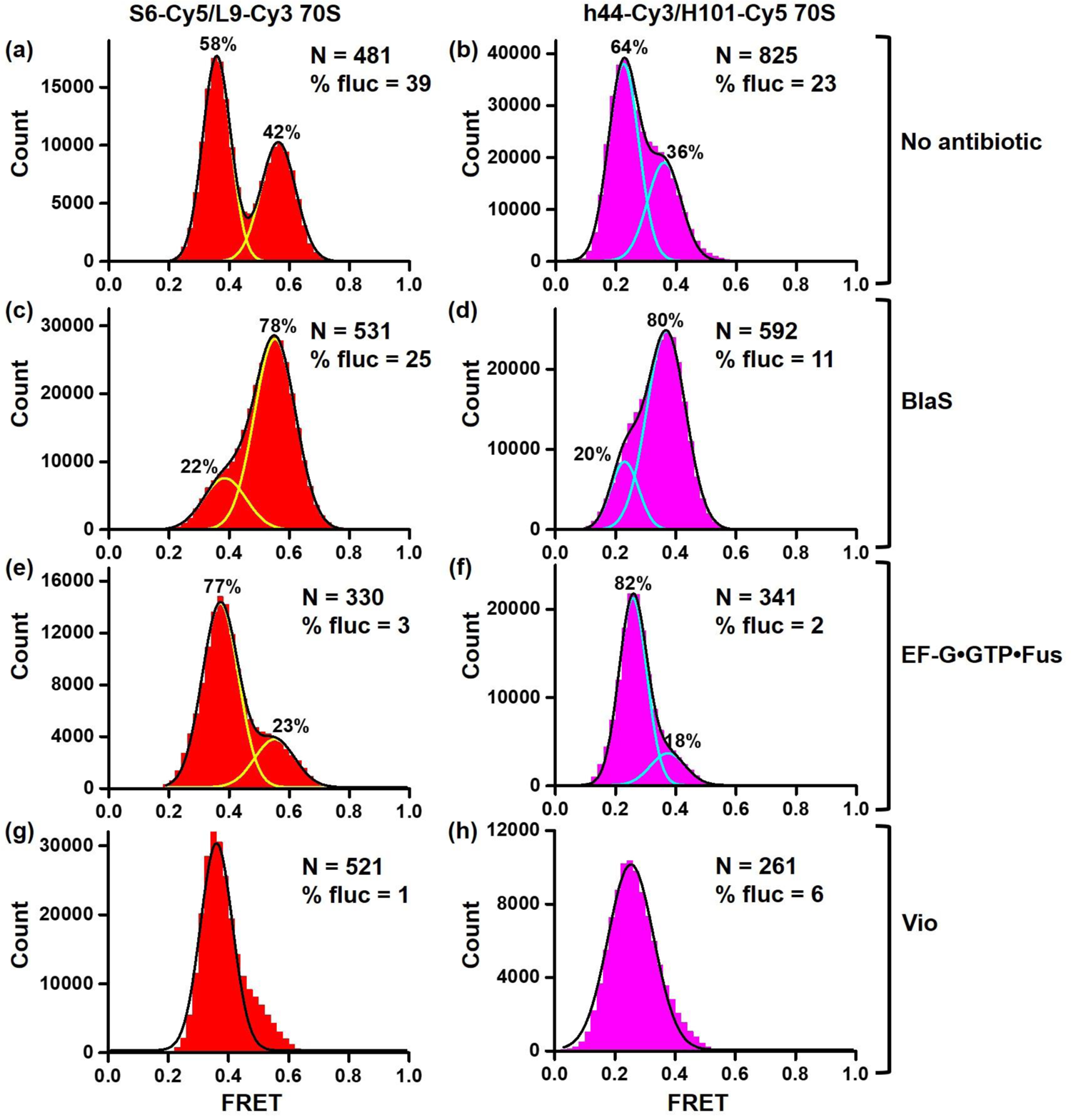
S6-Cy5/L9-Cy3 and h44-Cy3/H101-Cy5 assays reflect changes in the NR-R equilibrium. Histograms show FRET distribution in S6-Cy5/L9-Cy3 (a, c, e, g) and h44-Cy3/H101-Cy5 (b, d, f, h) ribosomes bound with P-site deacylated tRNA^fMet^ that were imaged in polyamine buffer in absence of antibiotics (a-b), in presence of BlaS (c-d), EF-G•GTP•Fusidic acid (e-f) or viomycin (g-h). N indicates the total number of FRET traces compiled into each histogram. % fluc designates the percentage of traces that show fluctuations. Cyan and yellow lines show individual Gaussian fits. Black line indicates the sum of Gaussian fits. The fraction of the ribosomes in the R and NR conformations are shown above the corresponding Gaussian peaks.

During translation initiation mediated by initiation factors IF1, IF2•GTP and IF3, ribosomal subunits form the post-initiation complex with the mRNA and initiator tRNA, fMet-tRNA^fMet^. The post-initiation ribosome bound with fMet-tRNA^fMet^ adopts the NR conformation [28]. S6-Cy5/L9-Cy3 and h44-Cy3/H101-Cy5 ribosomes were assembled with IF1, IF2•GTP, IF3, mRNA and fMet-tRNA^fMet^ in polyamine buffer (please see Materials and Methods), in which spontaneous intersubunit rotation was originally observed [12]. Consistent with structural data showing the post-initiation ribosome in the NR conformation [28], predominant 0.6 and 0.4 FRET states were observed in S6-Cy5/L9-Cy3 and h44-Cy3/H101-Cy5 post-initiation ribosomes, respectively (Fig. 1a-b). Only 10% of S6-Cy5/L9-Cy3 and h44-Cy3/H101-Cy5 ribosomes showed apparent FRET transitions between the NR and R conformations. The fraction of post-initiation ribosomes observed in the R conformation (31% of S6-Cy5/L9-Cy3 and 15% of h44-Cy3/H101-Cy5 ribosomes) might correspond to complexes in which fMet-tRNA^fMet^ was spontaneously deacylated and shifted to the hybrid state.

Published structural [28, 29] and FRET experiments [13] indicated that preventing IF2 release by replacing GTP with a non-hydrolysable analogue of GTP, GDPCP, traps the ribosome in the intermediate state of initiation, in which ribosomal subunits are rotated by 3-4° relative to each other. To test whether both intersubunit FRET pairs detect this semi-rotated conformation of the ribosome, we repeated subunit association in the presence of IF1, IF2•GDPCP and IF3. Consistent with published data, ∼0.5 FRET value was predominately observed in S6-Cy5/L9-Cy3 ribosomes that was halfway between 0.6 and 0.4 FRET and thus likely corresponded to the semi-rotated conformation of the ribosome (Fig. 1c). Similarly, ∼0.3 FRET value was predominately observed in h44-Cy3/H101-Cy5 ribosomes that was halfway between 0.4 and 0.2 FRET (Fig. 1d). These experiments indicated that both FRET assays detect the semi-rotated conformation of the ribosome sampled during translation initiation.

To mimic transition from the initiation phase of translation to the first elongation cycle, post-initiation ribosomes, containing fMet-tRNA^fMet^, were assembled with IF1, IF2•GTP and IF3, and were subsequently treated with antibiotic puromycin. Puromycin binds to the A site of the ribosome and acts as a substrate of peptidyl-transfer reaction effectively deacylating fMet-tRNA^fMet^. Deacylation of P-site tRNA triggers intersubunit rotation and tRNA movement into the hybrid state. Consistent with these expectations, treating S6-Cy5/L9-Cy3 and h44-Cy3/H101-Cy5 ribosomes with puromycin increased the presence of 0.4 and 0.2 FRET states, respectively, indicating that NR↔R equilibrium shifted to the R conformation (Fig. 1e-f). Taken together, our data show that under conditions known to stabilize the NR, R and semi-R conformations of the ribosome, the high, low and intermediate FRET values were predominantly observed in both S6-Cy5/L9-Cy3 and h44-Cy3/H101-Cy5 FRET pairs. Hence, both S6-Cy5/L9-Cy3 and h44-Cy3/H101-Cy5 FRET signals correlate with intersubunit rotation of the ribosome.

### S6/L9 and h44/H101 FRET pairs reveal spontaneous intersubunit rotation

Surprisingly, an appreciable number of h44-Cy3/H101-Cy5 ribosomes showed apparent spontaneous FRET fluctuations in the absence of EF-G (Fig. 1f). To further explore dynamics of h44-Cy3/H101-Cy5 ribosomes, post-initiation ribosomes containing fMet-tRNA^fMet^ in the P site were first incubated with EF-Tu•GTP•Phe-tRNA^Phe^ and then treated with EF-G•GTP to complete one elongation cycle. EF-Tu-mediated binding of aminoacyl-tRNA to the A site followed by peptidyl transfer from P- to A-site tRNA triggered switching from 0.4 (NR) to 0.2 (R) FRET in the majority of ribosomes (Fig. 2a). This is consistent with the formation of rotated, hybrid state conformation of the ribosome bound with A/P fMetPhe-tRNA^Phe^ and P/E deacylated tRNA^fMet^. Importantly, 16% of pre-translocation h44-Cy3/H101-Cy5 ribosomes imaged before addition of EF-G showed spontaneous transitions between 0.2 and 0.4 FRET states that are consistent with repetitive fluctuations between the non-rotated, classical-state and rotated, hybrid-state conformations of the ribosome (Fig. 2b). Catalyzed by EF-G•GTP, mRNA/tRNA translocation converted the majority of ribosomes from R (0.2 FRET) to NR (0.4 FRET) conformation (Fig. 2c).

To further explore dynamics of pre-translocation ribosomes containing deacylated tRNA in the P site, we non-enzymatically bound deacylated tRNA^fMet^ to the P site of either S6-Cy5/L9-Cy3 or h44-Cy3/H101-Cy5 ribosomes. Previous studies demonstrated that ribosomes containing deacylated tRNA in the P site show similar rates of spontaneous intersubunit rotation regardless of presence or absence of A-site peptidyl tRNA [12]. However, translocation does not occur in the absence of the A-site tRNA [30]. That makes ribosome complexes containing a single deacylated P-site tRNA a convenient model system to study dynamics of pre-translocation ribosomes due to absence of spontaneous translocation.

Both FRET pairs showed spontaneous fluctuations between the NR and R conformations in ribosomes containing P-site deacylated tRNA^fMet^ (Fig. 3a-b, Fig 4a-b). Ribosomes were not uniformly dynamic as spontaneous fluctuations were observed in only 39% of S6-Cy5/L9-Cy3 and 23% of h44-Cy3/H101-Cy5 (Fig. 3a-b). Interestingly, fluctuating S6-Cy5/L9-Cy3 and h44-Cy3/H101-Cy5 ribosomes typically displayed multiple FRET transitions per trace (Fig. 4). However, the equilibrium between the NR and R conformations was similarly shifted toward the R conformations in both fluctuating and non-fluctuating populations of S6-Cy5/L9-Cy3 and h44-Cy3/H101-Cy5 ribosomes (Supplementary Fig. 1). The rates of spontaneous NR→R transitions observed in S6-Cy5/L9-Cy3 and h44-Cy3/H101-Cy5 ribosomes were essentially identical (0.5 s^-1^). The rate of spontaneous R→NR transitions observed in S6-Cy5/L9-Cy3 ribosomes (0.3 s^-1^) was ∼2.5-fold slower than that in h44-Cy3/H101-Cy5 ribosomes (0.8 s^-1^) (Fig. 5 and Table 1).

**Figure 4.**
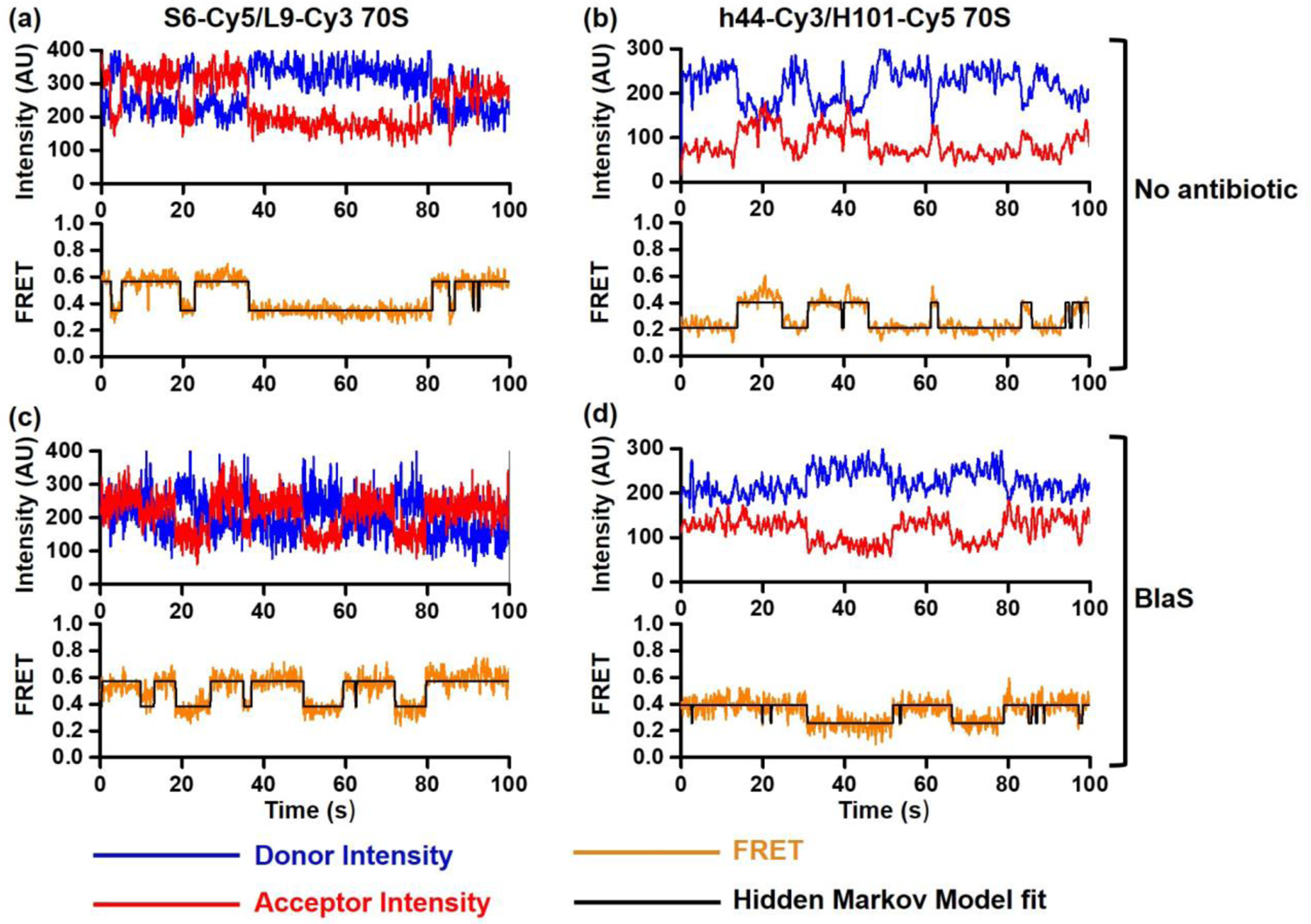
Spontaneous intersubunit rotation in S6-Cy5/L9-Cy3 and h44-Cy3/H101-Cy5 ribosomes imaged in polyamine buffer. smFRET traces show spontaneous intersubunit rotation in S6-Cy5/L9-Cy3 (a, c) and h44-Cy3/H101-Cy5 (b, d) ribosomes bound with P-site tRNA^fMet^ that were imaged in polyamine buffer in the absence (a-b) or presence of BlaS (c-d). Fluorescence intensities of Cy3 (donor) and Cy5 (acceptor) are shown in blue and red, respectively. FRET efficiency and HMM fit are shown in orange and black, respectively.

**Figure 5.**
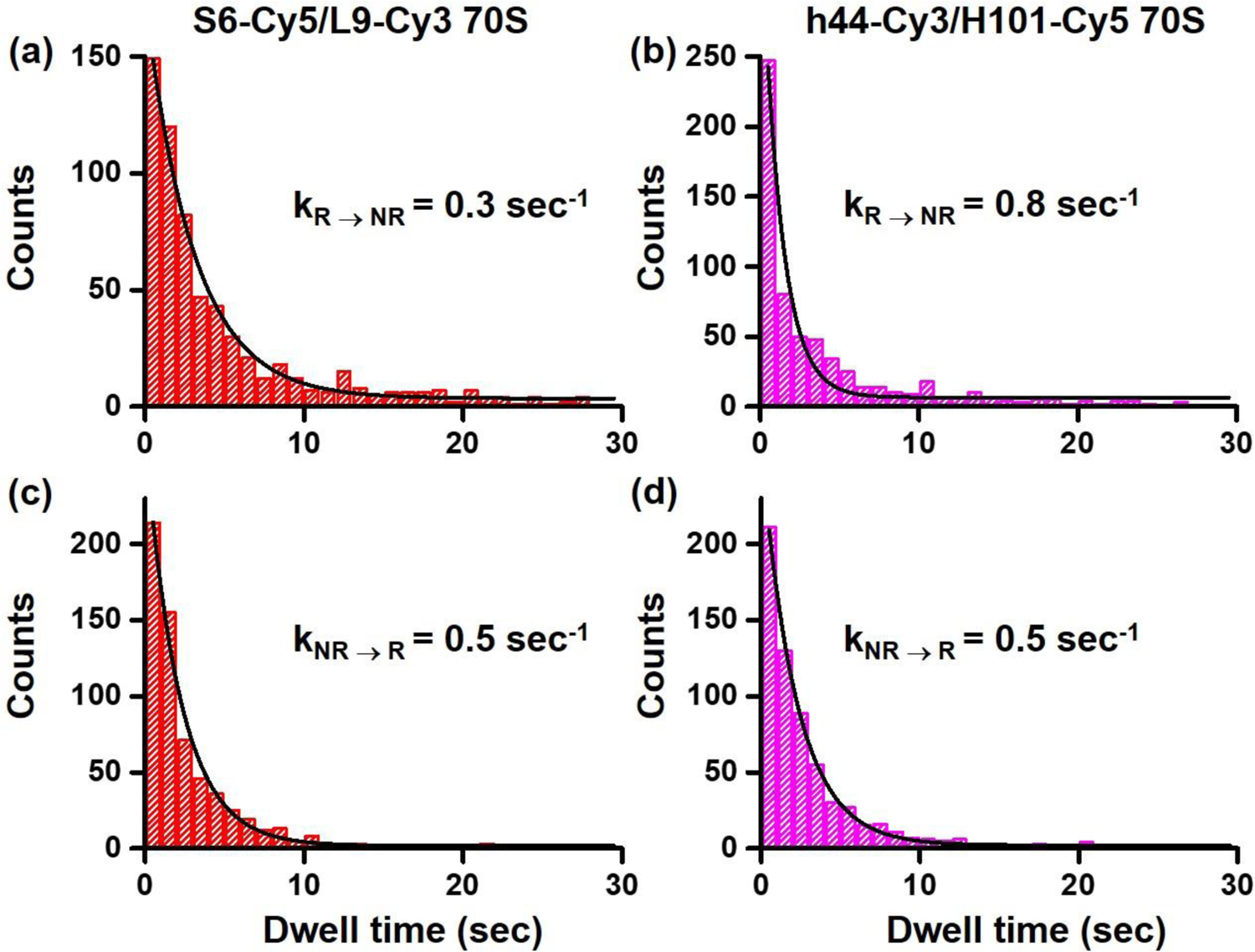
Rates of spontaneous intersubunit rotation in S6-Cy5/L9-Cy3 and h44-Cy3/H101-Cy5 ribosomes. Distributions of dwell times (1 second binning size) in the R (a-b) and NR (c-d) conformations in S6-Cy5/L9-Cy3 (a, c) and h44-Cy3/H101-Cy5 (b, d) ribosomes bound with P-site deacylated tRNA^fMet^ were fitted to single exponential decay (black line).

**Table 1.**
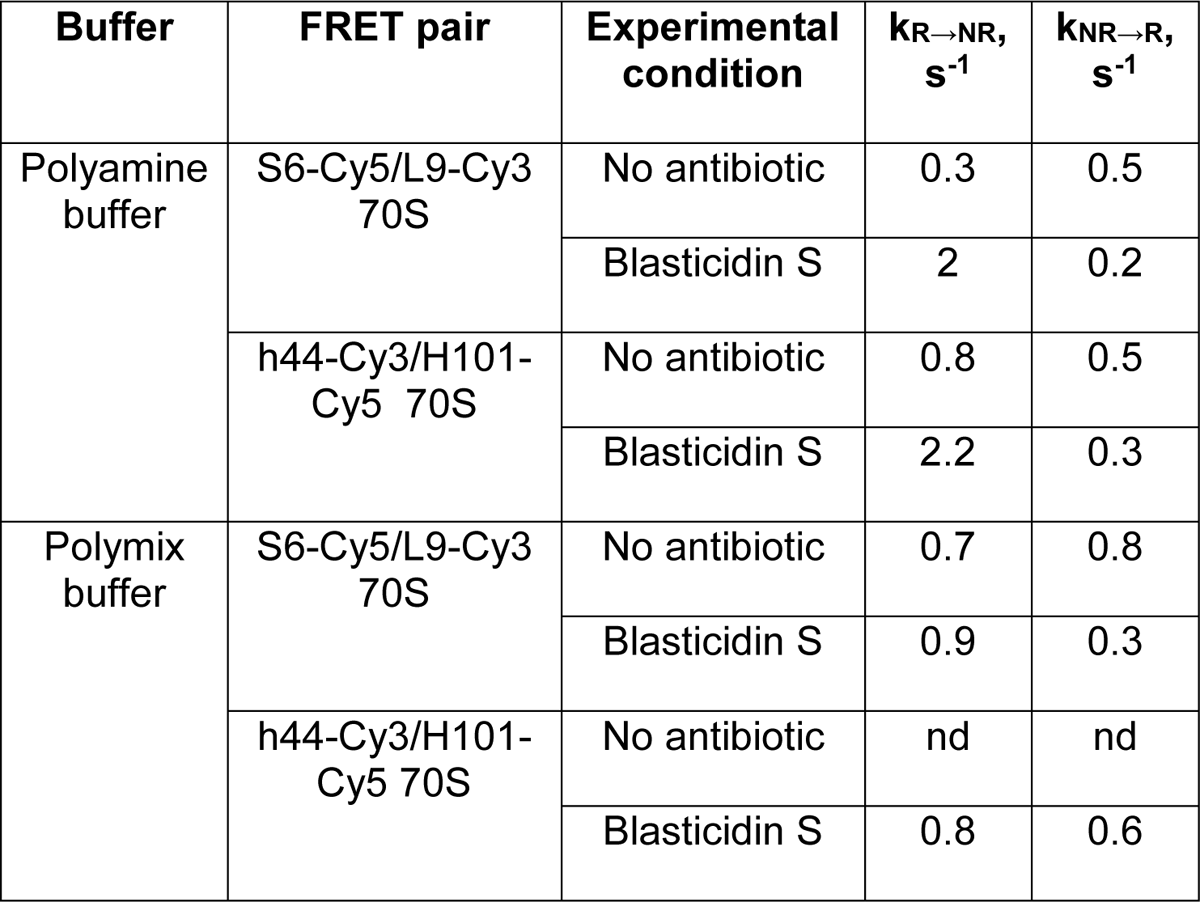
Rates of transitions between the NR and R conformations. Rates of transitions between the NR and R conformations in S6-Cy5/L9-Cy3 and h44-Cy3/H101-Cy5 ribosomes bound with P-site deacylated tRNA^fMet^ that were imaged in absence of antibiotics or in presence of BlaS. Rates were determined by fitting dwell time distributions (1 second binning size) to single exponential decay function. nd – not determined due lack of fluctuations.

Next, we tested whether translation factors and antibiotics, which were shown to perturb NR↔R equilibrium, affect S6-Cy5/L9-Cy3 and h44-Cy3/H101-Cy5 ribosomes differentially or similarly. Antibiotic Blasticidin S (BlaS) stabilizes the NR, classical state conformation of the ribosome by enhancing binding of deacylated tRNA to the 50S P site [31]. Consistent with this report, BlaS increased the fraction of the NR conformation from ∼40 to 80% in both S6-Cy5/L9-Cy3 and h44-Cy3/H101-Cy5 ribosomes (Fig. 3 c-d, Supplementary Fig. 2). Correspondingly, BlaS increased the rate of R→NR transition and caused reciprocal decrease in the rate of NR→R transition (Fig. 4 c-d, Table 1). Noteworthy, the rates of R→NR and NR→R transitions in BlaS-bound S6-Cy5/L9-Cy3 and h44-Cy3/H101-Cy5 ribosomes were remarkably similar (Table 1).

Binding of EF-G and antibiotic viomycin were previously shown to stabilize the rotated, hybrid state conformation of the ribosome [1, 12, 32]. When ribosomes containing deacylated P-site tRNA^fMet^ were incubated with EF-G•GTP and antibiotic fusidic acid, which inhibits EF-G disassociation after GTP hydrolysis, the fraction of the R conformation increased from 60 to 80% in both S6-Cy5/L9-Cy3 and h44-Cy3/H101-Cy5 ribosomes (Fig. 3 e-f). Notably, spontaneous intersubunit rotation nearly ceased and was observed in only 2 (h44-Cy3/H101-Cy5) – 3% (S6-Cy5/L9-Cy3) of ribosomes (Fig. 3 e-f, Table 1). Similarly, viomycin binding nearly completely converted h44-Cy3/H101-Cy5 and S6-Cy5/L9-Cy3 ribosomes into the R conformation and dramatically diminished the number of traces showing spontaneous FRET fluctuations (Fig. 3 g-h, Table 1).

Taken together, our data demonstrate that h44-Cy3/H101-Cy5 and S6-Cy5/L9-Cy3 FRET pairs respond similarly to changes in NR↔R equilibrium and exhibit similar rates of spontaneous NR→R and R→NR transitions. However, in the absence of EF-G or antibiotics as well as in the presence of BlaS, twice as many S6-Cy5/L9-Cy3 ribosomes show spontaneous intersubunit rotation than h44-Cy3/H101-Cy5 ribosomes (Fig. 3 a-d, Table 1).

### Spontaneous intersubunit dynamics is reduced in polymix buffer

We next tested how changes in buffer conditions affect intersubunit dynamics of h44-Cy3/H101-Cy5 and S6-Cy5/L9-Cy3 ribosomes. Aforementioned experiments were performed in polyamine buffer (please see Materials and Methods), which was originally used in studies demonstrating spontaneous intersubunit rotation in S6-Cy5/L9-Cy3 ribosomes [12]. By contrast, the original h44-Cy3/H101-Cy5 experiments were performed in the polymix buffer. The most significant difference between these two buffer systems seems to be concentration and identity of polyamines: polyamine buffer contains 2 mM spermidine and 0.1 mM spermine while polymix buffer contains 1 mM spermidine and 5 mM putrescine.

To examine effects of buffer conditions on intersubunit dynamics, we measured smFRET in pre-translocation ribosomes bound with P-site deacylated tRNA^fMet^ in polymix buffer. In both h44-Cy3/H101-Cy5 and S6-Cy5/L9-Cy3 ribosomes, replacing polyamine with polymix buffer did not significantly shift the NR↔R equilibrium. However, the fraction of ribosomes showing spontaneous fluctuations was reduced by 2 (S6-Cy5/L9-Cy3) and 20 folds (h44-Cy3/H101-Cy5), respectively. In fact, only 1% of h44-Cy3/H101-Cy5 ribosomes containing a deacylated P-site tRNA showed fluctuations between 0.2 and 0.4 FRET states that is consistent with published measurements of h44-Cy3/H101-Cy5 smFRET in polymix buffer [11].

When antibiotic BlaS was incubated with ribosomes in polymix buffer, nearly 90% of both h44-Cy3/H101-Cy5 and S6-Cy5/L9-Cy3 ribosomes were observed in the NR conformation consistent with ability of BlaS to stabilize the NR, classical state conformation of the ribosome. Addition of BlaS did not substantially affected the fraction of S6-Cy5/L9-Cy3 ribosomes showing spontaneous fluctuations. By contrast, BlaS produced ∼10-fold increase in the fraction of h44-Cy3/H101-Cy5 ribosomes showing spontaneous fluctuations. In agreement with experiments performed in the polyamine buffer, the rates of spontaneous R→NR transitions observed in BlaS-bound S6-Cy5/L9-Cy3 and h44-Cy3/H101-Cy5 ribosomes were similar (0.9 s^-1^ and 0.8 s^-1^, respectively). The rate of spontaneous NR→R transitions observed in BlaS-bound S6-Cy5/L9-Cy3 ribosomes (0.3 s^-1^) was two-fold slower than that in h44-Cy3/H101-Cy5 ribosomes (0.6 s^-1^).

When either EF-G•GTP•fusicid acid or viomycin were added, the R conformation was predominantly observed in both S6-Cy5/L9-Cy3 and h44-Cy3/H101-Cy5 ribosomes. Only 5% or less of S6-Cy5/L9-Cy3 and h44-Cy3/H101-Cy5 ribosomes bound with EF-G or viomycin showed spontaneous fluctuations between the NR and R conformations. These results agree with published data [12] and our experiments performed in the polyamine buffer that demonstrate stabilization of the R conformation by EF-G and viomycin (Fig. 3, Table 1).

Our results show that h44-Cy3/H101-Cy5 and S6-Cy5/L9-Cy3 FRET pairs respond similarly to changes in NR↔R equilibrium in both polyamine and polymix buffers. However, polymix buffer significantly reduces the fraction of ribosomes exhibiting FRET fluctuations in S6-Cy5/L9-Cy3 and especially h44-Cy3/H101-Cy5 assays. These observations, at least partially, reconcile discrepancies between results previously obtained using S6-Cy5/L9-Cy3 assay in polyamine buffer and h44-Cy3/H101-Cy5 assay in polymix buffer.

## Discussion

Our side-by-side comparison of S6-Cy5/L9-Cy3 and h44-Cy3/H101-Cy5 FRET pairs performed in two different buffer systems reaffirms that both FRET assays reliably report changes in the equilibrium between the NR and R conformations of the ribosomes and thus can be used to follow intersubunit rotation during translation. In accord with this conclusion, both FRET assays produced self-consistent results in numerous published studies [11-15, 33-35] while the S6/L9 assay was also employed by several independent research groups [12, 13, 36–38].

Both S6-Cy5/L9-Cy3 and h44-Cy3/H101-Cy5 assays reveal spontaneous intersubunit rotation in pre-translocation ribosomes containing deacylated P-site tRNA demonstrating that GTP hydrolysis by EF-G is not required for the transition from the R into NR conformation. Interestingly, in both S6-Cy5/L9-Cy3 and h44-Cy3/H101-Cy5 constructs, fluctuating traces typically show multiple FRET transitions per trace (Fig. 4 and 7). Remarkably, fluctuating S6-Cy5/L9-Cy3 and h44-Cy3/H101-Cy5 ribosomes exhibit similar kinetics of spontaneous NR→R and R→NR transitions (Table 1). The observed rates of R→NR transitions (0.3 - 2 s^-1^) are only marginally slower than the rates of EF-G-catalyzed translocation measured at the same experimental conditions (i.e. room temperature and buffer system) [10, 39, 40]. These results are consistent with the idea that the large-scale rearrangements of the ribosome associated with translation are predominately thermally driven and do not require GTP-hydrolysis by factors of translation.

Our experiments demonstrate that rather than being an invariable inherent property of each FRET construct, the fraction of ribosomes exhibiting FRET fluctuations changes depending on experimental conditions such as buffer system and the presence of antibiotics. Hence, non-fluctuating ribosomes are not irreversibly damaged or inactive. At least to some extent, variations in the fraction of fluctuating ribosomes likely reflect changes in the rates of NR→R and R→NR transitions that shift the NR↔R equilibrium. Because of limited time resolution (100 ms here and in most published studies) and Cy5 photobleaching in smFRET experiments, very fast and very slow fluctuations evade detection giving rise to seemingly static traces. Changes in the rates of NR→R and R→NR transitions favoring either the NR or R conformations can increase the fraction of apparently static traces. In agreement with this idea, stabilization of the R, hybrid state conformation by viomycin, or the presence of peptidyl-tRNA in the P site, which shifts the NR↔R equilibrium toward the NR, classical state conformation, significantly diminish the fraction of spontaneously fluctuating ribosomes [12] (Fig. 3, 6). Likewise, variations in the propensity of deacylated P-site tRNA to adopt the hybrid (P/E) state were demonstrated to change the fraction of S6-Cy5/L9-Cy3 ribosomes exhibiting spontaneous fluctuations from ∼10 (P-site tRNA^Tyr^) to 70% (P-site tRNA^fMet^) [12]. Similarly, smFRET between fluorophores attached to the elbow of P-site tRNA and protein uL1 of the L1 stalk also showed that the fraction of pre-translocation ribosomes, which displayed spontaneous tRNA fluctuations between the classical and hybrid states, varied between ∼20 and 80% depending on identity of deacylated P-site tRNA [19].

Shifts in the NR↔R equilibrium do not fully account for observed differences in the fraction of fluctuating ribosomes. For example, in both S6-Cy5/L9-Cy3 and h44-Cy3/H101-Cy5 ribosomes, switching from polyamine to polymix buffer greatly reduced the number of traces displaying apparent FRET transitions without altering the NR↔R equilibrium (Fig. 3 and 6). In addition, FRET distribution histograms compiled from traces lacking FRET transitions were generally similar to histograms compiled from traces showing spontaneous fluctuations (Supplementary Fig. 1-2). Hence, the fraction of molecules displaying apparent fluctuations might be influenced by the presence of subpopulations of ribosomes with different kinetics of intersubunit rotation. This kinetic heterogeneity might be due to a reversible conformational rearrangement such as disruption of one of intersubunit bridges that might produce faster-fluctuating ribosomes. Extensions of helixes h44 and H101 for fluorescent labeling might alter the frequency of the hypothesized switching from static to dynamic subpopulations and thus diminish the fraction of ribosomes showing FRET fluctuations.

**Figure 6.**
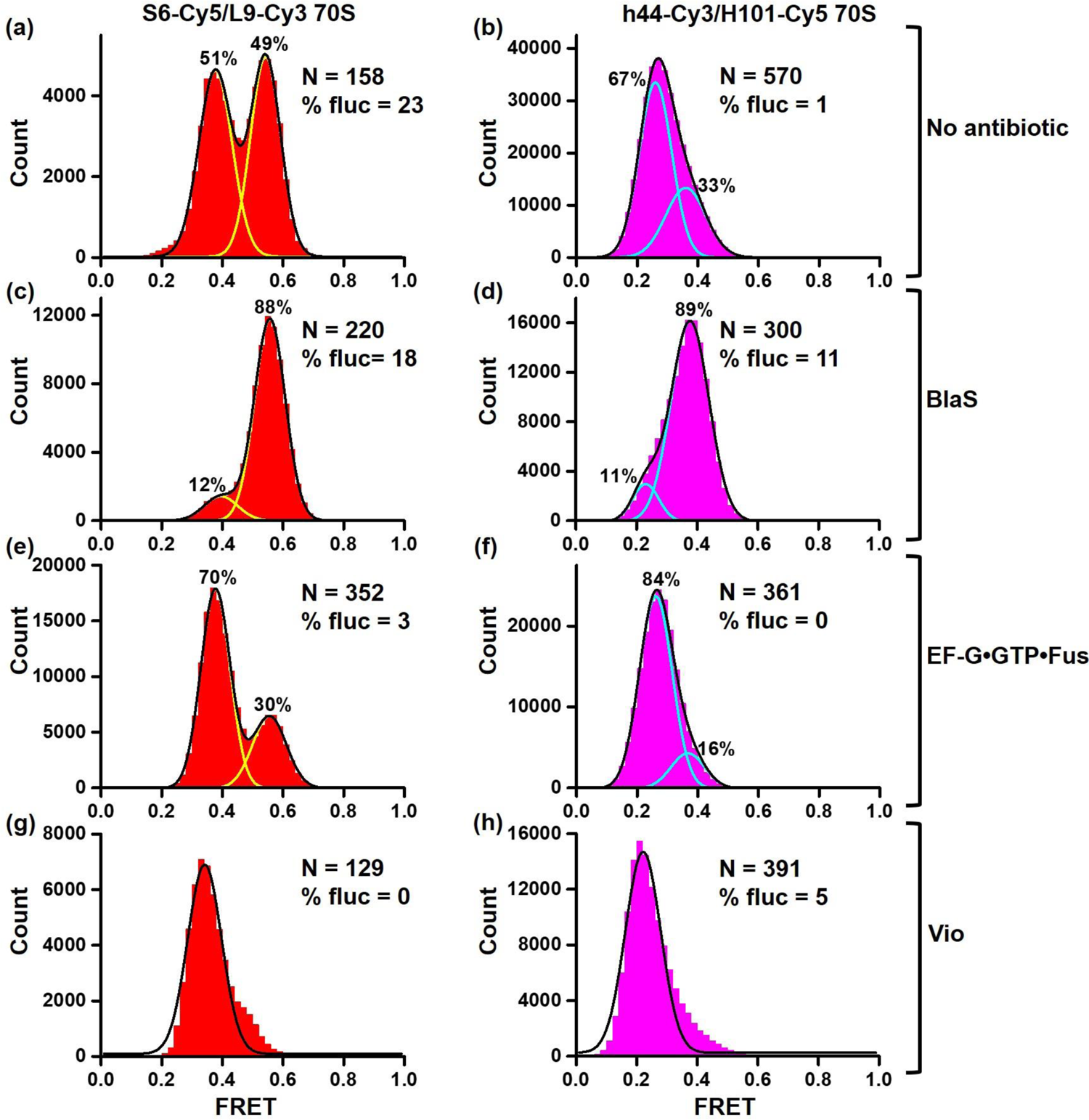
Following changes in the NR-R equilibrium in polymix buffer system. Histograms show FRET distribution in S6-Cy5/L9-Cy3 (a, c, e, g) and h44-Cy3/H101-Cy5 (b, d, f, h) ribosomes bound with P-site deacylated tRNA^fMet^ that were imaged in polymix buffer in absence of antibiotics (a-b), in presence of BlaS (c-d), EF-G•GTP•Fusidic acid (e-f) or viomycin (g-h). N indicates the total number of FRET traces compiled into each histogram. % fluc designates the percentage of traces that show fluctuations. Cyan and yellow lines show individual Gaussian fits. Black line indicates the sum of Gaussian fits. The fraction of the ribosomes in the R and NR conformations are shown above the corresponding Gaussian peaks.

**Figure 7.**
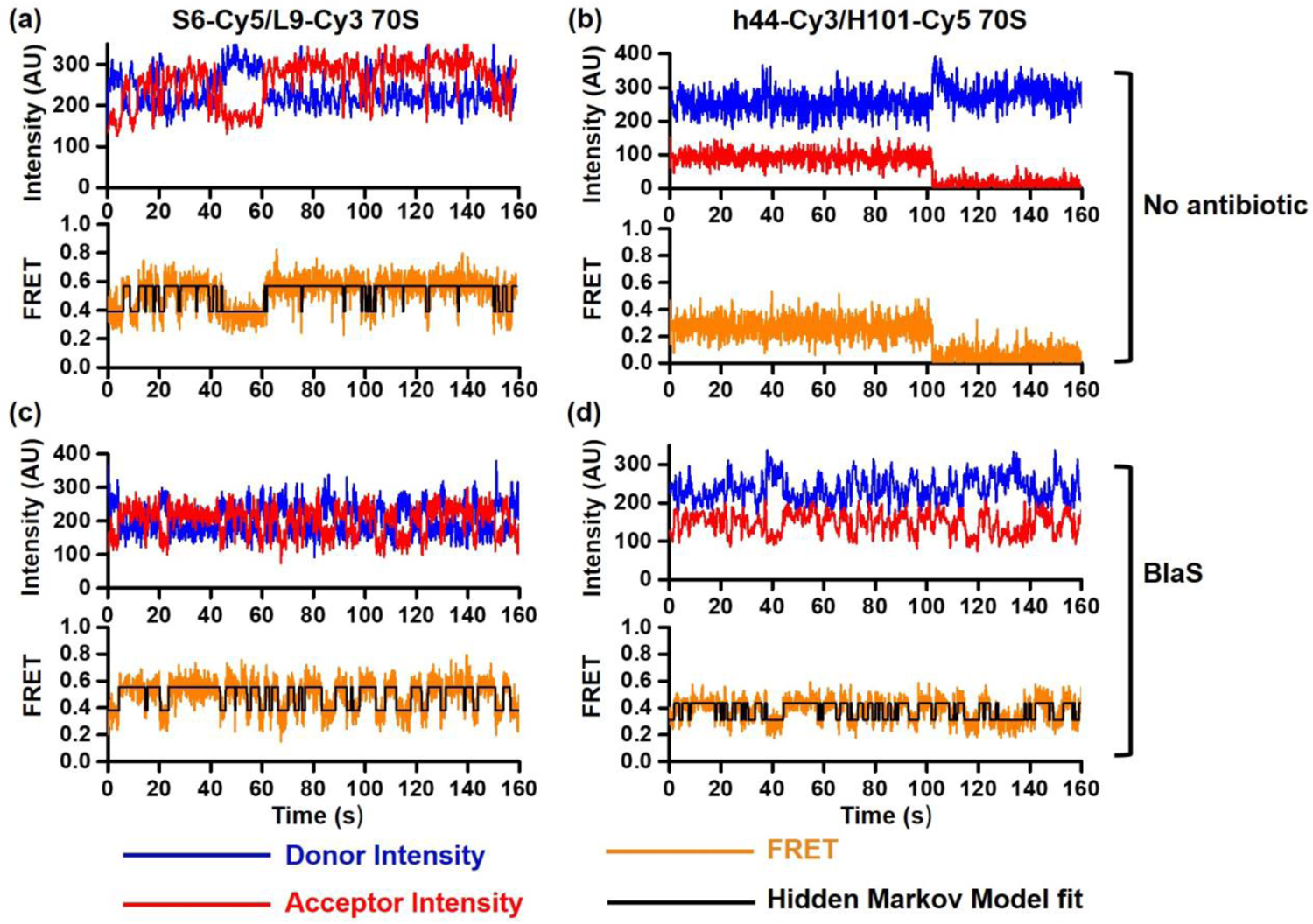
Spontaneous intersubunit rotation in S6-Cy5/L9-Cy3 and h44-Cy3/H101-Cy5 ribosomes imaged in polymix buffer system. smFRET traces show spontaneous intersubunit rotation in S6-Cy5/L9-Cy3 (a, c) and h44-Cy3/H101-Cy5 (b, d) ribosomes bound with P-site tRNA^fMet^ that were imaged in polymix buffer in the absence (a-b) or presence of BlaS (c-d). Fluorescence intensities of Cy3 (donor) and Cy5 (acceptor) are shown in blue and red, respectively. FRET efficiency and HMM fit are shown in orange and black, respectively.

Finally, molecular dynamics simulations revealed that with some frequency, the intrinsic structural flexibility of the ribosomal subunits might leave the distance between fluorophores in h44-Cy3/H101-Cy5 pair nearly unchanged despite intersubunit rotation [27, 41]. Experimental conditions such as concentrations of polyamines might also influence the intrinsic structural flexibility of the ribosomal subunits that interferes with FRET detection of intersubunit rotation.

While we cannot unequivocally explain the differences in behavior of S6-Cy5/L9-Cy3 and h44-Cy3/H101-Cy5 FRET pairs, our results highlight importance of using multiple FRET pairs especially in studies of large, complex and dynamic macromolecular assemblies.

## Materials and Methods

### Buffer systems

Experiments were performed in either in polyamine buffer (50 mM HEPES, pH 7.5, 6 mM MgCl_2_, 150 mM NH_4_Cl, 0.1 mM spermine, 2 mM spermidine and 6 mM β-mercaptoethanol) or polymix buffer (50 mM Tris-OAc, pH 7.5, 5 mM MgCl_2_, 100 mM KCl, 5 mM NH_4_Ac, 0.5 mM CaCl_2_, 5 mM putrescine, 1 mM spermidine, 0.5 mM EDTA) as indicated.

### Preparation of ribosome, translation factors, tRNA and mRNA

**Cy3 and Cy5** maleimide were purchased from Click Chemistry Tools. S6-Cy5/L9-Cy3 ribosomes were prepared by partial reconstitution of ΔS6-30S and ΔL9-50S subunits with S6-41Cys-Cy5 and L11-11Cys-Cy3 proteins as previously described [13, 39]. The 30S and 50S ribosomal subunits containing rRNA extensions for hybridization with fluorescent oligos in h44 and H101, respectively, were provided by Alexey Petrov (Auburn University). To label h44/H101 of 70S ribosomes 1.5 µM of H101 50S was incubated with 3 µM of sp22-Cy5 DNA oligo [25] at 37°C for 10 minutes and then another 10 minutes at 30°C. Likewise, 1 µM of h44 30 S was incubated with 2 µM of sp68-Cy3 oligo [25]. Labelled 50S and labelled 30S were then mixed and incubated at 37°C for 10 minutes. tRNA^fMet^ and tRNA^Phe^ (purchased from Chemical Block) were aminoacylated using S100 cell extract; Met-tRNA^fMet^ was formylated as previously described [9]. Histidine-tagged EF-Tu and EF-G were expressed and purified by standard procedures.

### Ribosome complex assembly

To assemble P-site tRNA ribosome complexes, 0.3 μM S6/L9-labeled (or h44/H101-labeled) 70S ribosomes were incubated with 0.6 μM tRNA^fMet^ and 0.6 μM HIV_NS ΔFSS mRNA for at 37°C 10 minutes. HIV_ΔFSS mRNA which was pre-annealed to 3’ biotin-AL2 DNA oligo (HIV_NS ΔFSS mRNA: 5’ *GGUUUUUCUUCUGAAGAUAAAG*CAACAACAACAAGGCA**AGGAGG**UAAAA<u>AUG</u>UU CUACAA 3’; AL2-complementray sequence is italicized; Shine-Dalgarno sequence is in **bold**; AUG codon is <u>underlined</u>). To assemble initiation complexes (IC), 0.3 μM 30S was incubated with 0.6 μM fMet-tRNA^fMet^, 0.6 μM HIV_NS ΔFSS mRNA (pre-annealed to 3’ biotin-AL2 oligo), 0.6 μM IF1, 0.6 μM IF2, 0.6 μM IF3 and 1 mM GTP (or GDPCP) at 37°C for 10 minutes. Next, 0.15 μM 30S IC and 0.24 μM 50S subunits were associated at 37°C for 10 minutes. To bind aminoacyl-tRNA to the A site of post-initiation complex, 0.3 μM Phe-tRNA^Phe^ was bound with 5 μM EF-Tu and 1 mM GTP at 37°C for 5 minutes. Then 0.15 μM 70S was incubated with 0.3 μM Phe-tRNA^Phe^•EF-Tu•GTP at 37°C for 10 minutes. Translocation was catalyzed by 2 μM EF-G and 0.5 mM GTP. Ribosomal complexes were diluted to 2 nM before smFRET imaging.

### smFRET measurements

Quartz slides were coated with dichlorodimethylsilane (DDS), bound with biotin-BSA and passivated by Tween-20. To enable tethrering to biotin-BSA, 30 μL 0.2 mg/mL neutravidin dissolved in H50 buffer (20 mM HEPES, pH 7.5 and 50 mM KCl). Non-specific sample binding to the slide was checked in the absence of neutravidin. Ribosomal complexes were imaged in polyamine buffer or polymix buffer containing 0.8 mg/mL glucose oxidase, 0.625% glucose, 0.4 μg/mL catalase and 1.5 mM 6-hydroxy-2,5,7,8-tetramethylchromane-2-carboxylic acid (Trolox). 1 mM blasticidin S (Fisher Scientific), 0.5 mM viomycin (USP), 2 µM EF-G, 0.5 mM GTP (Sigma-Aldrich), 1 mM puromycin (Sigma-Aldrich) or 40 µM fusidic acid (Sigma-Aldrich) were added into imaging buffer as indicated. smFRET was excited by 532 nm laser and measured by prism-type inverted TIRF (Total Internal Reflection Fluorescence) microscope with a water-immersion objective (Olympus, UplanApo, 60x, NA 1.2). A DV2 (Photometrics) dual-view imaging system equipped with a 630-nm dichroic mirror was used to split the fluorescence into Cy3 and Cy5 channels. TIRF microscopy signal has been acquired by “Single” software http://ha.med.jhmi.edu/resources/) and detected with an EMCCD camera (Andor iXon) with 100 ms integration time and with 300 EM gain.

IDL software (ITT) was used to extract flourescence intensities of Cy3 donor (I *_D_*) and Cy5 acceptor (I*_A_*), from which apparent FRET efficiency (E *_FRET_*, hence referred as FRET) was calculated:

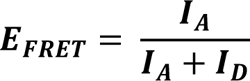

Traces with single-step photobleachings for both Cy5 and Cy3 were selected using MATLAB. FRET distribution histograms compiled from hundreds of smFRET traces were smoothed with a 5-point window using MATLAB and fit to two Gaussians corresponding to 0.36 (±0.02) and 0.56 (±0.02) for S6-Cy5/L9-Cy3 ribosomes; 0.2 (±0.02) and 0.35 (±0.01) for h44-Cy3/H101-Cy5 ribosomes. FRET distribution histograms show number of data points (frames) vs FRET (0.02 binning size). To determine the rate of fluctuation smFRET traces were idealized by 2-state Hidden Markov Model (HMM) using HaMMy software (http://ha.med.jhmi.edu/resources/) [42]. To test whether the number of transitions in h44-Cy3/H101-Cy5 smFRET traces was underestimated by HaMMy, we re-analyzed S6-Cy5/L9-Cy3 and h44-Cy3/H101-Cy5 smFRET traces using another HMM algorithm, ebFRET [43, 44]. HaMMy and ebFRET detected similar number of transitions in both S6-Cy5/L9-Cy3 and h44-Cy3/H101-Cy5 smFRET traces (Supplementary Table 1).

## ACKNOWLEDGEMENTS

The 30S and 50S ribosomal subunits containing rRNA extensions for hybridization with fluorescent oligos in h44 and H101, respectively, were gifts from Alexey Petrov. We thank Alexey Petrov and Paul Whitford for helpful discussions. This work was supported by grant of the National Institutes of Health R35GM141812 (D.N.E.).

## AUTHORS CONTRIBUTIONS

D.N.E. designed the project. A.D. and C.B. performed experiments. A.D. and D.N.E. wrote the manuscript.

## References

[1] Frank J, Agrawal RK. A ratchet-like inter-subunit reorganization of the ribosome during translocation. Nature. 2000;406:318–22.

[2] Frank J, Gonzalez RL, Jr. Structure and dynamics of a processive Brownian motor: the translating ribosome. Annu Rev Biochem. 2010;79:381–412.

[3] Moazed D, Noller HF. Intermediate states in the movement of transfer RNA in the ribosome. Nature. 1989;342:142–8.

[4] Valle M, Zavialov A, Sengupta J, Rawat U, Ehrenberg M, Frank J. Locking and unlocking of ribosomal motions. Cell. 2003;114:123–34.

[5] Agirrezabala X, Lei J, Brunelle JL, Ortiz-Meoz RF, Green R, Frank J. Visualization of the hybrid state of tRNA binding promoted by spontaneous ratcheting of the ribosome. Mol Cell. 2008;32:190–7.

[6] Julian P, Konevega AL, Scheres SH, Lazaro M, Gil D, Wintermeyer W, et al. Structure of ratcheted ribosomes with tRNAs in hybrid states. Proc Natl Acad Sci U S A. 2008;105:16924–7.

[7] Noller HF, Lancaster L, Zhou J, Mohan S. The ribosome moves: RNA mechanics and translocation. Nat Struct Mol Biol. 2017;24:1021–7.

[8] Korostelev AA. The Structural Dynamics of Translation. Annu Rev Biochem. 2022;91:245–67.

[9] Ermolenko DN, Majumdar ZK, Hickerson RP, Spiegel PC, Clegg RM, Noller HF. Observation of intersubunit movement of the ribosome in solution using FRET. J Mol Biol. 2007;370:530–40.

[10] Ermolenko DN, Noller HF. mRNA translocation occurs during the second step of ribosomal intersubunit rotation. Nat Struct Mol Biol. 2011;18:457–62.

[11] Marshall RA, Dorywalska M, Puglisi JD. Irreversible chemical steps control intersubunit dynamics during translation. Proc Natl Acad Sci U S A. 2008;105:15364–9.

[12] Cornish PV, Ermolenko DN, Noller HF, Ha T. Spontaneous intersubunit rotation in single ribosomes. Mol Cell. 2008;30:578–88.

[13] Ling C, Ermolenko DN. Initiation factor 2 stabilizes the ribosome in a semirotated conformation. Proc Natl Acad Sci U S A. 2015;112:15874–9.

[14] Aitken CE, Puglisi JD. Following the intersubunit conformation of the ribosome during translation in real time. Nat Struct Mol Biol. 2010;17:793–800.

[15] Bao C, Loerch S, Ling C, Korostelev AA, Grigorieff N, Ermolenko DN. mRNA stem-loops can pause the ribosome by hindering A-site tRNA binding. Elife. 2020;9.

[16] Blanchard SC, Kim HD, Gonzalez RL, Jr., Puglisi JD, Chu S. tRNA dynamics on the ribosome during translation. Proc Natl Acad Sci U S A. 2004;101:12893–8.

[17] Munro JB, Altman RB, O’Connor N, Blanchard SC. Identification of two distinct hybrid state intermediates on the ribosome. Mol Cell. 2007;25:505–17.

[18] Fei J, Kosuri P, MacDougall DD, Gonzalez RL, Jr. Coupling of ribosomal L1 stalk and tRNA dynamics during translation elongation. Mol Cell. 2008;30:348–59.

[19] Fei J, Richard AC, Bronson JE, Gonzalez RL, Jr. Transfer RNA-mediated regulation of ribosome dynamics during protein synthesis. Nat Struct Mol Biol. 2011;18:1043–51.

[20] Cornish PV, Ermolenko DN, Staple DW, Hoang L, Hickerson RP, Noller HF, et al. Following movement of the L1 stalk between three functional states in single ribosomes. Proc Natl Acad Sci U S A. 2009;106:2571–6.

[21] Fei J, Bronson JE, Hofman JM, Srinivas RL, Wiggins CH, Gonzalez RL, Jr. Allosteric collaboration between elongation factor G and the ribosomal L1 stalk directs tRNA movements during translation. Proc Natl Acad Sci U S A. 2009;106:15702–7.

[22] Munro JB, Altman RB, Tung CS, Cate JH, Sanbonmatsu KY, Blanchard SC. Spontaneous formation of the unlocked state of the ribosome is a multistep process. Proc Natl Acad Sci U S A. 2010;107:709–14.

[23] Ling C, Ermolenko DN. Structural insights into ribosome translocation. Wiley Interdiscip Rev RNA. 2016;7:620–36.

[24] Spirin AS. The ribosome as a conveying thermal ratchet machine. J Biol Chem. 2009;284:21103–19.

[25] Dorywalska M, Blanchard SC, Gonzalez RL, Kim HD, Chu S, Puglisi JD. Site-specific labeling of the ribosome for single-molecule spectroscopy. Nucleic Acids Res. 2005;33:182–9.

[26] Chen J, Tsai A, O’Leary SE, Petrov A, Puglisi JD. Unraveling the dynamics of ribosome translocation. Curr Opin Struct Biol. 2012;22:804–14.

[27] Levi M, Nguyen K, Dukaye L, Whitford PC. Quantifying the Relationship between Single-Molecule Probes and Subunit Rotation in the Ribosome. Biophys J. 2017;113:2777–86.

[28] Kaledhonkar S, Fu Z, Caban K, Li W, Chen B, Sun M, et al. Late steps in bacterial translation initiation visualized using time-resolved cryo-EM. Nature. 2019;570:400–4.

[29] Sprink T, Ramrath DJ, Yamamoto H, Yamamoto K, Loerke J, Ismer J, et al. Structures of ribosome-bound initiation factor 2 reveal the mechanism of subunit association. Sci Adv. 2016;2:e1501502.

[30] Joseph S, Noller HF. EF-G-catalyzed translocation of anticodon stem-loop analogs of transfer RNA in the ribosome. EMBO J. 1998;17:3478–83.

[31] Svidritskiy E, Ling C, Ermolenko DN, Korostelev AA. Blasticidin S inhibits translation by trapping deformed tRNA on the ribosome. Proc Natl Acad Sci U S A. 2013;110:12283–8.

[32] Ermolenko DN, Spiegel PC, Majumdar ZK, Hickerson RP, Clegg RM, Noller HF. The antibiotic viomycin traps the ribosome in an intermediate state of translocation. Nat Struct Mol Biol. 2007;14:493–7.

[33] Marshall RA, Aitken CE, Puglisi JD. GTP hydrolysis by IF2 guides progression of the ribosome into elongation. Mol Cell. 2009;35:37–47.

[34] Chen J, Petrov A, Tsai A, O’Leary SE, Puglisi JD. Coordinated conformational and compositional dynamics drive ribosome translocation. Nat Struct Mol Biol. 2013;20:718–27.

[35] Uemura S, Aitken CE, Korlach J, Flusberg BA, Turner SW, Puglisi JD. Real-time tRNA transit on single translating ribosomes at codon resolution. Nature. 2010;464:1012–7.

[36] Desai BJ, Gonzalez RL, Jr. Multiplexed genomic encoding of non-canonical amino acids for labeling large complexes. Nat Chem Biol. 2020;16:1129–35.

[37] Qin P, Yu D, Zuo X, Cornish PV. Structured mRNA induces the ribosome into a hyper-rotated state. EMBO Rep. 2014;15:185–90.

[38] Sharma H, Adio S, Senyushkina T, Belardinelli R, Peske F, Rodnina MV. Kinetics of Spontaneous and EF-G-Accelerated Rotation of Ribosomal Subunits. Cell Rep. 2016;16:2187–96.

[39] Rexroad G, Donohue JP, Lancaster L, Noller HF. The role of GTP hydrolysis by EF-G in ribosomal translocation. Proc Natl Acad Sci U S A. 2022;119:e2212502119.

[40] Salsi E, Farah E, Ermolenko DN. EF-G Activation by Phosphate Analogs. J Mol Biol. 2016;428:2248–58.

[41] Hassan A, Byju S, Freitas FC, Roc C, Pender N, Nguyen K, et al. Ratchet, swivel, tilt and roll: a complete description of subunit rotation in the ribosome. Nucleic Acids Res. 2022.

[42] McKinney SA, Joo C, Ha T. Analysis of single-molecule FRET trajectories using hidden Markov modeling. Biophys J. 2006;91:1941–51.

[43] van de Meent JW, Bronson JE, Wood F, Gonzalez RL, Jr., Wiggins CH. Hierarchically-coupled hidden Markov models for learning kinetic rates from single-molecule data. JMLR Workshop Conf Proc. 2013;28:361–9.

[44] van de Meent JW, Bronson JE, Wiggins CH, Gonzalez RL, Jr. Empirical Bayes methods enable advanced population-level analyses of single-molecule FRET experiments. Biophys J. 2014;106:1327–37.

