## Supplemental Table 1, Supplemental Figures 1 and 2 for "Comparing FRET pairs that report on intersubunit rotation in bacterial ribosomes"

**Supplementary Figure 1**

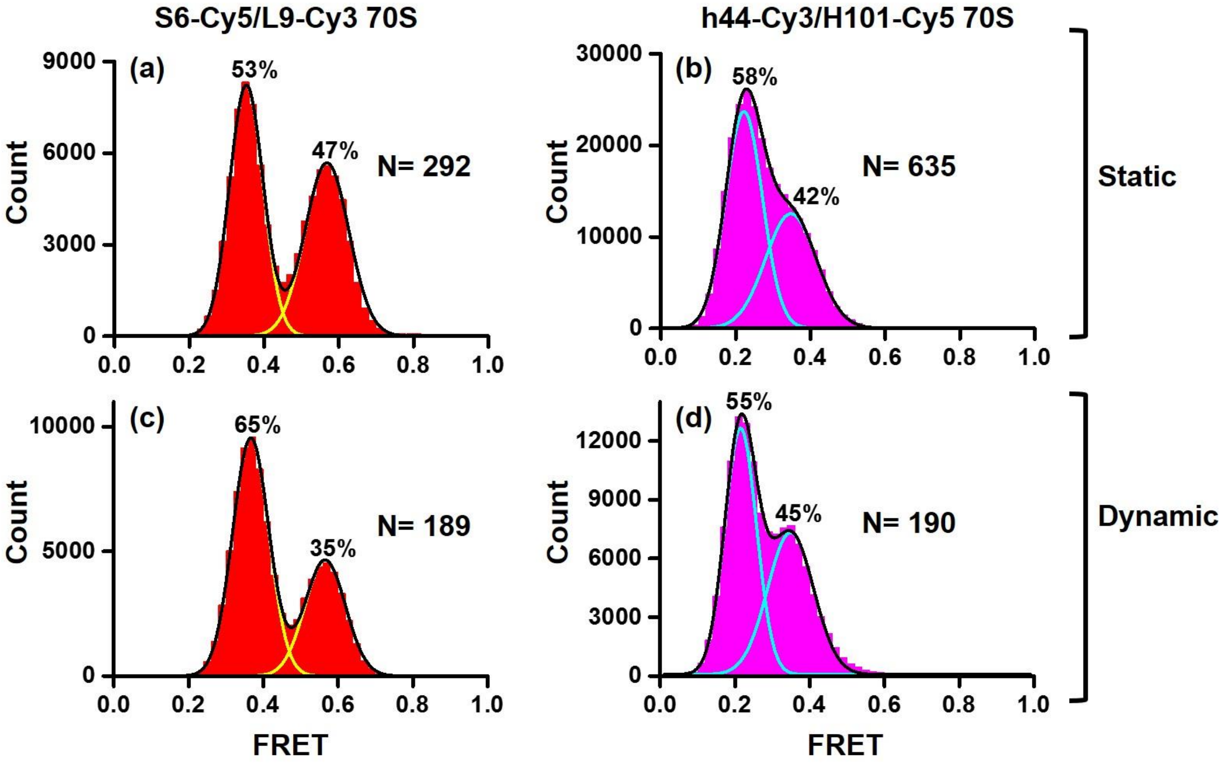


**Static and dynamic traces exhibit similar FRET distribution.** Histograms show FRET distribution in S6-Cy5/L9-Cy3 (a, c) and h44-Cy3/H101-Cy5 (b, d) ribosomes bound with deacylated P-site $tRNA^{fMet}$ that were imaged in polyamine buffer in absence of antibiotics. Histograms were compiled from either “static” traces (a-b), which lack FRET transitions, or “dynamic” traces (c-d) that show spontaneous FRET fluctuations. N indicates the total number of FRET traces. Yellow and cyan lines show individual Gaussian fits of FRET distribution. Black line indicates the sum of Gaussian fits. Corresponding FRET histograms that combine static and dynamic traces are shown in Fig. 3a (S6-Cy5/L9-Cy3) and Fig. 3b (h44-Cy3/H101-Cy5).

**Supplementary Figure 2**

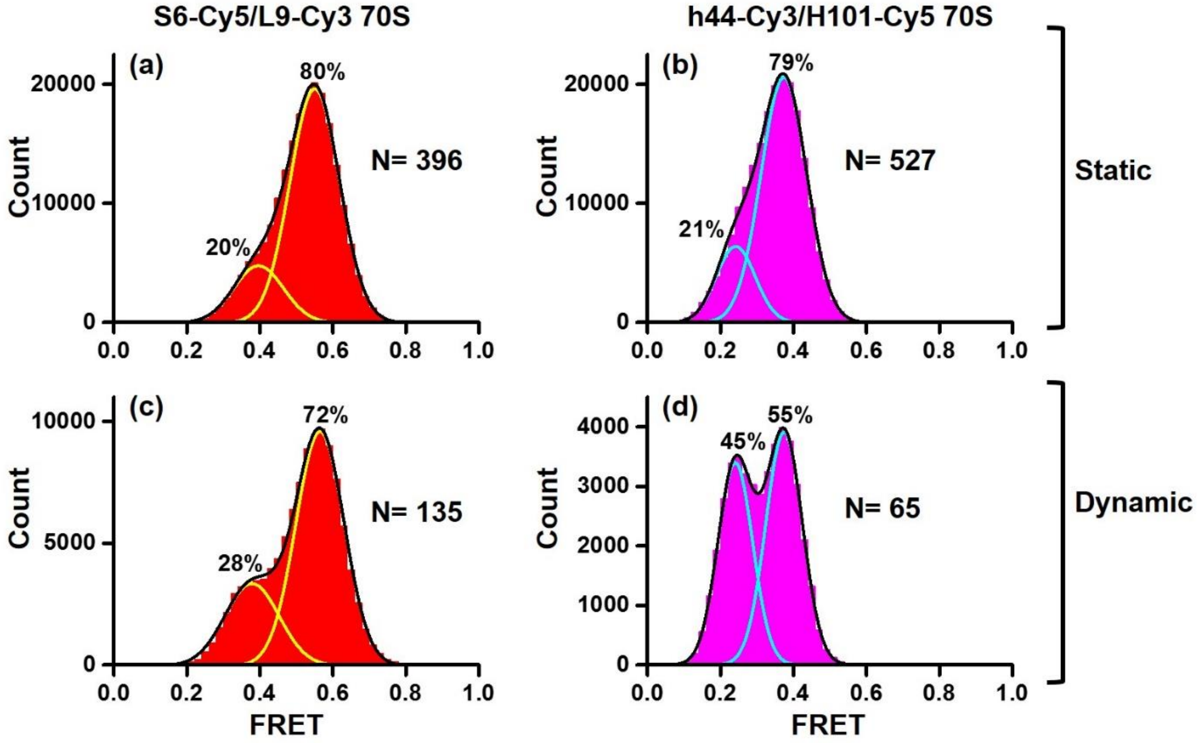


**In BlaS-bound ribosomes, static and dynamic traces exhibit similar FRET distribution.** Histograms show FRET distribution in S6-Cy5/L9-Cy3 (a, c) and h44-Cy3/H101-Cy5 (b, d) ribosomes bound with deacylated P-site tRNA$^{fMet}$ that were imaged in polyamine buffer in the presence of BlaS. Histograms were compiled from either static traces (a-b), which lack FRET transitions, or dynamic traces (c-d) that show spontaneous FRET fluctuations. N indicates the total number of FRET traces. Yellow and cyan lines show individual Gaussian fits of FRET distribution. Black line indicates the sum of Gaussian fits. Corresponding FRET histograms that combine static and dynamic traces are shown in Fig. 3c (S6-Cy5/L9-Cy3) and Fig. 3d (h44-Cy3/H101-Cy5).

**Supplementary Table 1. Number of FRET transitions detected by HaMMy and ebFRET algorithms in S6-Cy5/L9-Cy3 and h44-Cy3/H101-Cy5 smFRET traces.**

| Buffer | FRET pair | Experimental condition | Number of R→NR transitions | | Number of NR→R transitions | |
|---|---|---|---|---|---|---|
| | | | **HaMMy** | **ebFRET** | **HaMMy** | **ebFRET** |
| Polyamine buffer | S6-Cy5/L9-Cy3 70S | No antibiotic | 643 | 657 | 630 | 644 |
| | | Blasticidin S | 462 | 444 | 452 | 429 |
| | h44-Cy3/H101-Cy5 70S | No antibiotic | 643 | 652 | 637 | 637 |
| | | Blasticidin S | 391 | 311 | 365 | 291 |
| Polymix buffer | S6-Cy5/L9-Cy3 70S | No antibiotic | 284 | 284 | 276 | 273 |
| | | Blasticidin S | 336 | 357 | 365 | 352 |
| | h44-Cy3/H101-Cy5 70S | No antibiotic | nd | nd | nd | nd |
| | | Blasticidin S | 372 | 376 | 362 | 367 |

Transitions between the NR and R conformations in S6-Cy5/L9-Cy3 and h44-Cy3/H101-Cy5 ribosomes bound with deacylated P-site $tRNA^{fMet}$ that were imaged in absence of antibiotics or in presence of BlaS were detected by fitting smFRET traces using HaMMy and ebFRET HMM algorithms.
